# Computational analysis on the ACE2-derived peptides for neutralizing the ACE2 binding to the spike protein of SARS-CoV-2

**DOI:** 10.1101/2020.05.03.075473

**Authors:** Cecylia S. Lupala, Vikash Kumar, Xuanxuan Li, Xiao-dong Su, Haiguang Liu

**Author notes:** These authors contributed equally (Cecylia S. Lupala, Vikash Kumar). Corresponding author: Haiguang Liu.

## Abstract

The severe acute respiratory syndrome coronavirus 2 (SARS-CoV-2), the causative agent of the COVID-19, is spreading globally and has infected more than 3 million people. It has been discovered that SARS-CoV-2 initiates the entry into cells by binding to human angiotensin-converting enzyme 2 (hACE2) through the receptor binding domain (RBD) of its spike glycoprotein. Hence, drugs that can interfere the SARS-CoV-2-RBD binding to hACE2 potentially can inhibit SARS-CoV-2 from entering human cells. Here, based on the N-terminal helix α1 of human ACE2, we designed nine short peptides that have potential to inhibit SARS-CoV-2 binding. Molecular dynamics simulations of peptides in the their free and SARS-CoV-2 RBD-bound forms allow us to identify fragments that are stable in water and have strong binding affinity to the SARS-CoV-2 spike proteins. The important interactions between peptides and RBD are highlighted to provide guidance for the design of peptidomimetics against the SARS-CoV-2.

## Introduction

Severe acute respiratory syndrome coronavirus 2 (SARS-CoV-2, also known as 2019-nCoV) caused the COVID-19, which has been declared by the World Health Organization to be a global pandemic. The COVID-19 has caused over 219,000 fatalities (as of April 29th, 2020) with more than 3.1 million people testing positive for the coronavirus1. The virus causes influenza-like symptoms in patients with mild symptoms while severe cases are reported to develop severe lung injury that leads to multi-organ failures, eventually death^2–5^. The rapid growth of COVID-19 infections all over the world requires global efforts to fight against the virus^1,6^.

The phylogenetic analysis revealed that the SARS-CoV-2 belongs the genus betacoronavirus and possesses about 96% nucleotide sequence identity with the closest bat coronavirus RaTG13, which was identified in horseshoe bats (*Rhinolophus species*) in 2013. It shares 79% similarities with SARS-CoV genome, and its genome has 89% identity to two other bat SARS-like viruses (Bat-SL-CoVZC45 and Bat-SL-CoVZXC21)^7,8^. Both SARS-CoV and SARS-CoV-2 utilize the human angiotensin-converting enzyme 2 (hACE2) to initiate the spike protein binding and facilitate the viral attachment to host cells. *In vitro* and *in vivo* studies have confirmed hACE2 as the functional receptor of SARS-CoV^9–14^. For SARS-CoV, it has been shown that the overexpression of ACE2 enhances disease severity in mice infected with the virus. This revealed that ACE2-dependent entry of SARS-CoV into the host cells is a critical step^15^. Other studies on SARS-CoV have also reported that injecting SARS-CoV spike glycoproteins into mice decreased ACE2 expression levels and worsened lung injury^16,17^. Therefore ACE2 is critical as both the entry receptor of SARS-CoV and in SARS-CoV pathogenesis ACE2 protects the lungs from injury^13^. Given the close relation between the spike proteins from SARS-CoV and SARS-CoV-2, the roles of hACE2 are crucial in the virus infection. Without affecting the ACE2 expression levels, designing molecules that can interfere the virus binding to hACE2 is highly desirable in the fight against the COVID-19.

Structural studies of SARS-CoV-2 spike glycoprotein show that the spike protein directly binds to ACE2 and their binding affinity is higher than that of SARS-CoV^18,19^. Studies have further shown that the 193-residue RBD of the SARS-CoV or SARS-CoV-2 spike protein is sufficient to bind to the human ACE2^10,20^. Based on these facts, the RBD of SARS-CoV-2 is considered a critical protein model for drug development to treat the COVID-19. Recently, both computational and experimental studies have reported the usage of ACE2 proteins as a method to block SARS-CoV-2 entry^21–23^. Clinical-grade human-recombinant ACE2 was demonstrated to markedly inhibit SARS-CoV-2 infections of the infected vascular organoids. The study also showed human-recombinant ACE2 reduced SARS-CoV-2 recovery levels from Vero cells by a factor of >1000, demonstrated to be effective in blocking virus infections^22^. The spike protein binding to hACE2 was also investigated in another study, which aims to develop molecules that interfere the binding of SARS-CoV-2 RBD to hACE2^21^. Their results showed that a 23-residue peptide (residues 21-43) of hACE2 N-terminal helix was able to bind to the RBD with nanomolar affinity, comparable to that of full length hACE2. They also reported that a 12-residue peptide (residues 27-38) failed to bind to the SARS-CoV-2 RBD. In a computational study, a 31-residue peptide derived from hACE2 with residues 22-44 and 351-357 (another critical binding site for the RBD) linked via a glycine was designed. The peptide binding affinity was improved by optimizing the peptide sequences through sequence substitutions^23^. These studies demonstrate that recombinant ACE2 and short peptides derived from hACE2 can provide a line of defense against SARS-CoV-2 infection. Furthermore, studies on SARS-COV binding with hACE2 reported that hACE2 fragments composed of residues 22-44 or 22-57 inhibited the binding of SARS-CoV RBD to the human ACE2 with IC50 values of about 50 μM and 6 μM, respectively^24^, implying that the longer peptide fragment of residues 22-57 had stronger binding affinity to the SARS-CoV RBD, providing a more wiggle room for longer peptide design.

In the present work, we have designed nine peptides derived from the N-terminal helix of human ACE2, the α1 helix, with various lengths (from 12 to 70 residues), with an aim to maintain the direct interactions observed between the SARS-CoV-2 spike protein and hACE2 in the crystal structure ^11,20^. These peptides, or **S**pike **I**nteracting **F**ragments (SIFs), were modeled and simulated in water and in complex with the RBD of SARS-CoV-2 spike protein. The stability and binding affinities are quantified from the simulation data, providing molecular basis for the SIF design.

## Materials and methods

The crystal structure of SARS-CoV2-RBD/ACE2 complex (PDB ID: 6ZLG^11^) was used as the template to design peptides, which are subject to extensive MD simulations. Each SIF was simulated in a solvent box in its free form to study the peptide structural and dynamical stability in solvent. For those peptides exhibiting expected structure characteristics and high stability in solution, they are modelled in complex with SARS-CoV-2 RBD to further investigate the binding affinities. For all systems, parameterization and equilibration input files were prepared using the CHARMM-GUI webserver^25^. Each system was solvated in TIP3P water and sodium chloride ions to neutralize the systems to a salt concentration of 150 mM. The molecular systems were modeled with the CHARMM36 force field^26^.

After energy minimization using the steepest descent algorithm, each system was equilibrated at human body temperature 310.15 K, which was maintained by Nose-Hoover scheme with 1.0 ps coupling constant in the NVT ensemble (constant volume and temperature) for 125.0 ps under periodic boundary conditions with harmonic restraint forces applied to the complex molecules (400 kJ mol^−1^ nm^−2^ on backbone and 40 kJ mol^−1^ nm^−2^ on the side chains). In the subsequent step, the harmonic restraints forces were removed and the NPT ensemble were simulated at one atmosphere pressure (105 Pa) and body temperature. The pressure was maintained by isotropic Parrinello-Rahman barostat at 1.0 bar, with a compressibility of 4.5 × 10^−5^ bar^−1^ and coupling time constant of 5.0 ps. In all simulations, a time step of 2.0 fs was used and the PME (particle mesh Ewald) was applied for long-range electrostatic interactions. The van der Waals interactions were evaluated within the distance cutoff of 12.0 Å. Hydrogen atoms were constrained using the LINCS algorithm^27^. Each SIF peptide system was simulated for 300 ns and for the SARS-CoV2-RBD-SIF peptide complexes, simulation trajectories of 500 ns were propagated, using the GROMACS 5.1.2 package^28^.

Analyses were carried out with tools in the GROMACS (such as *rmsd*, *rmsf* and *do_dssp*) to examine the structural and dynamical properties, including the overall stability, residue and general structure fluctuations through the simulations. The VMD^29^ and Chimera^30^ software were used to analyze the hydrogen bonds, molecular binding interface, visualization, and to render images.

### MM-GBSA interaction energy calculation

Interaction energy calculation was carried out by Prime 3.0 MM-GBSA module of the SCHRODINGER^31^. To reduce uncertainties due to a single structure, 11 frames belonging to last 100ns MD simulations were used to calculate MM-GBSA interaction energies. In the case of SIF-RBD complex, SIF and RBD were considered as ligand and receptor respectively. Prime MM-GBSA uses OPLS-AA force field and VSGB 2.0 implicit solvation model to estimate the binding energy of the receptor-ligand complex. The binding energy is calculated as:

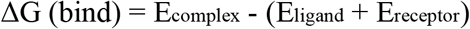

Where ΔG (bind) is the energy difference between the complex and sum of the energies of receptor and ligand alone. Energy for complex, receptor and ligand can be further divided into molecular mechanical and solvation (polar and non-polar) components.

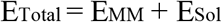

## Results

### Native interactions between SARS-CoV-2 RBD with human ACE2

Analysis of crystal structure of SARS-CoV-2 spike protein with human ACE2 revealed that the residues of the spike RBD makes extensive interactions with the N-terminal residues of hACE2 (19-83) (**Figure 1A**). The RBD-ACE2 interface contains a mixture of charged, polar and non-polar residues. We classified interactions into three types, hydrogen bonds, salt bridges and van der Waals (vdW) interactions. In the crystal structure, several hydrogen bonds and salt bridges exist at the interface between the RBD and hACE2. Apart from the hydrogen bonds, we also observed vdW contacts that contribute to the binding of RBD to hACE2. Major interactions listed in the **Table 1** were well maintained throughout MD simulations of the RBD/hACE2 complexes (**Figure 1B**).

**Figure 1.**
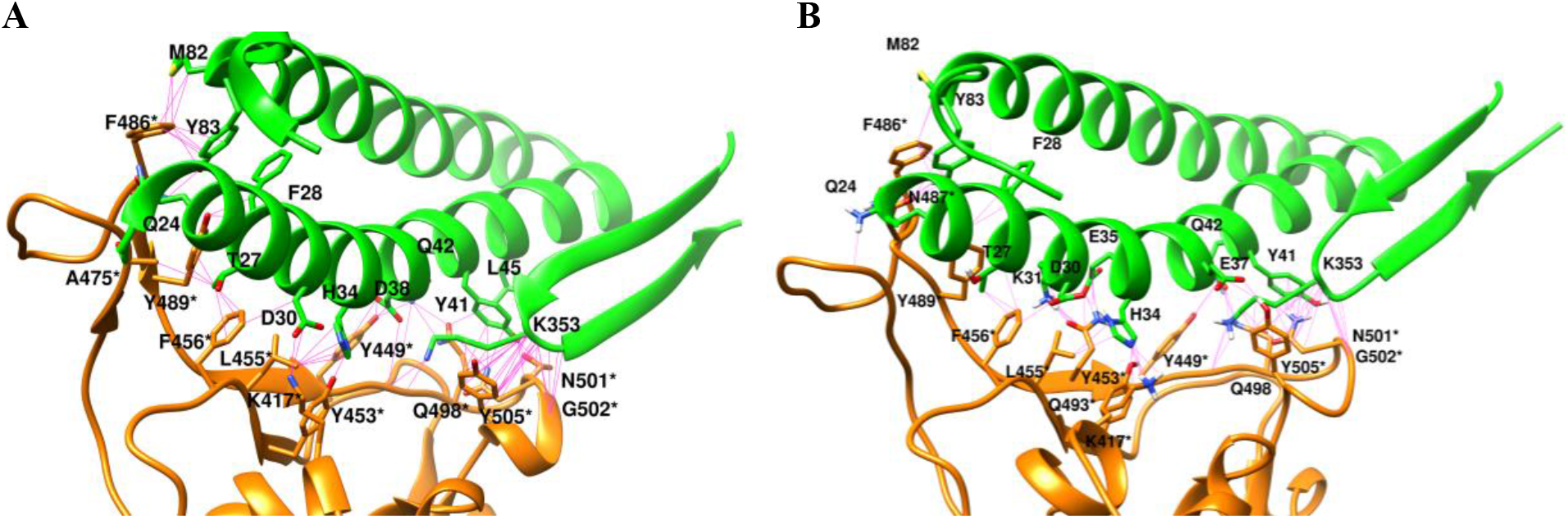
Interactions between SARS-CoV-2 RBD and hACE2. (A) The interactions revealed in the crystal structure. (B) The final structure of the complex in the 300 ns simulation. The van der Waals contacts between human ACE2 (green) and RBD (orange) of SARS-CoV-2 are shown with lines in magenta color. For clarity, only the direct interacting region of the hACE2 protein is shown.

**Table 1.**
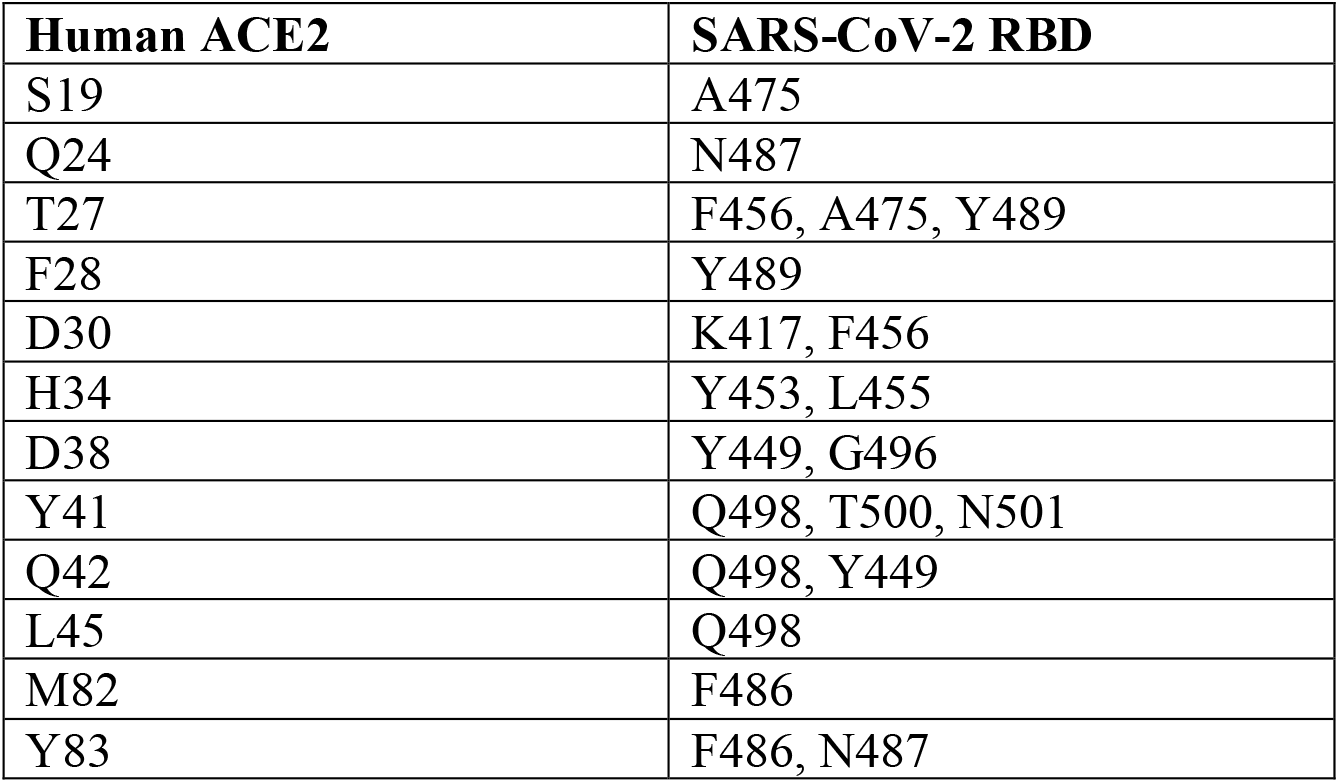
Contacting residues between hACE2 and SARS-CoV-2 RBD in the crystal structure (PDB: 6LZG).

### SARS-Cov-2 RBD interactions with the SIFs derived from hACE2

Based on the crystal structure and MD simulations of RBD-hACE2 complex, we designed the peptides that potentially bind to the SARS-CoV-2 spike protein by grafting the sequences from the N-terminal region of the hACE2. Several fragments with lengths ranging from 12 to 70 were selected for analysis (**Figure 2**). We analyzed all trajectories in terms of conformational changes, occupancies of H-bonds, and number of contacts between RBD and SIFs. Among 9 peptides, SIF5 and SIF8 have been reported in previous work^21^. SIF5 contains residues 21-43 of hACE2, while SIF8 is composed of hACE2 residues 27-38. The short SIF8 (12-residue) was reported to fail binding with the spike protein, while the SIF5 binds strongly.

**Figure 2.**
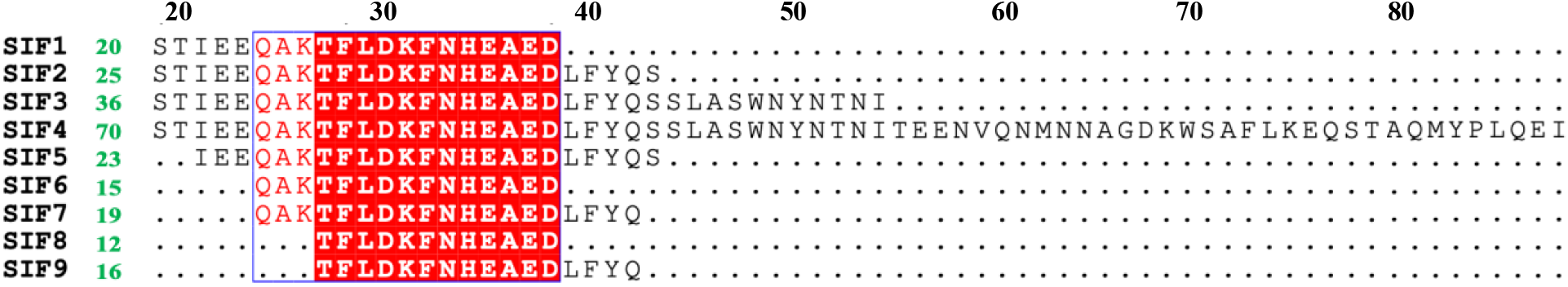
Detailed information about the hACE2 derived peptides. The lengths of the peptides are indicated to the right of their SIF IDs in green color.

### Short peptides lose helicity in the water

Structural stabilities of peptide fragments in the water were analyzed based on MD simulation results. Helical contents of all fragments are listed in **Table 2**. We observed that three peptide fragments, SIF3, SIF4 and SIF5 (residues 19-54, 19-88 and 21-43), maintained helicity higher than 75% in the water. Peptides composed of residues 19-38, 19-43 and 24-42 exhibited helicity between 60 and 70%. Interestingly, three short peptides 24-38, 27-38 and 27-42 lost more than 50% of the helical contents in the water, as shown in **Figure 3**. For all designed SIFs, their interactions with the spike RBD strongly depend on conformations, which were designed to be mostly helices as they are in the full hACE2 protein. The SIF’s with higher helical contents are stronger competitors with full hACE2 for spike protein binding (see the binding energy analysis), therefore they are more likely to be drug candidates for inhibiting the binding.

**Table 2.** Preliminary quantitative analysis of the MD Simulation trajectories.

| Peptides | RMSD of SIF alone in water | RMSD of SIF in complex with spike protein RBD | Helical content of SIF alone in water (%) | Helical content of SIF in complex with spike protein RBD (%) |
| --- | --- | --- | --- | --- |
| SIF1 | 5.03±1.53 | 3.45 ±0.55 | 62 | **79** |
| SIF2 | 4.20±2.09 | 4.85 ±1.92 | 68 | **80** |
| SIF3 | 2.36±0.68 | 5.62 ±1.68 | **90** | **79** |
| SIF4 | 2.48±0.59 | 3.19 ±1.03 | **83** | **82** |
| SIF5 | 4.47±2.07 | 4.21 ±1.30 | **80** | 62 |
| SIF6 | 4.69±1.19 | 3.21 ±0.99 | 31 | 63 |
| SIF7 | 3.17±1.74 | 4.96 ±2.00 | 62 | 67 |
| SIF8 | 4.43±1.17 | 6.96 ±2.60 | 45 | 52 |
| SIF9 | 3.94±2.05 | 8.92 ± 2.70 | 34 | 34 |

**Figure 3.**
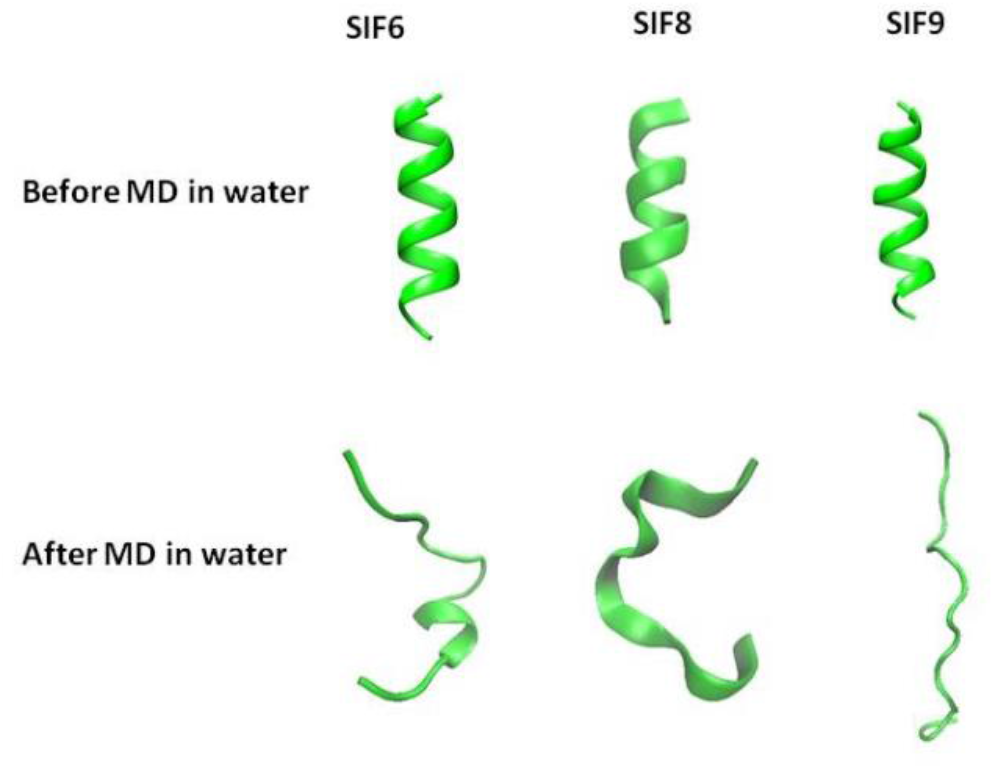
Short peptides lose their helicities in the water revealed by simulations.

The final conformations for the SIF/RBD complexes are shown in **Figure 4** for all nine designed peptides. Some peptides remained closely bound to the RBD, while some other peptides went through large conformational changes, ended with different structures from helices, which are the starting structures for these SIFs. We carried out quantitative analysis for these SIF’s, in both cases with and without the Spike RBD of SARS-CoV-2 (**Table 2**). The longest peptide (SIF4), which is 70-residue long and has 3 helices, showed an average RMSD 2.48±0.59Å in water, with respect to its conformation in the crystal structure of the full hACE2. In contrast, the peptide SIF-8 was found to be unstable is solvent with average RMSD of 4.43±1.17Å, indicating large deviation from its conformation in the full hACE2. Several SIF peptides (SIF3, SIF7, SIF8, and SIF9 in particular) exhibited larger conformational changes when they are bound to the RBD than in the free peptide forms. This observation suggests that those peptides prefer a similar helical conformation as they were in the full hACE2 protein in solution, and the binding to the RBD resulted induced conformational changes, which are more pronounced for shorter peptides, such as SIF8 and SIF9. Under the consideration of peptide stability, longer peptides are preferred according to the simulation results.

**Figure 4.**
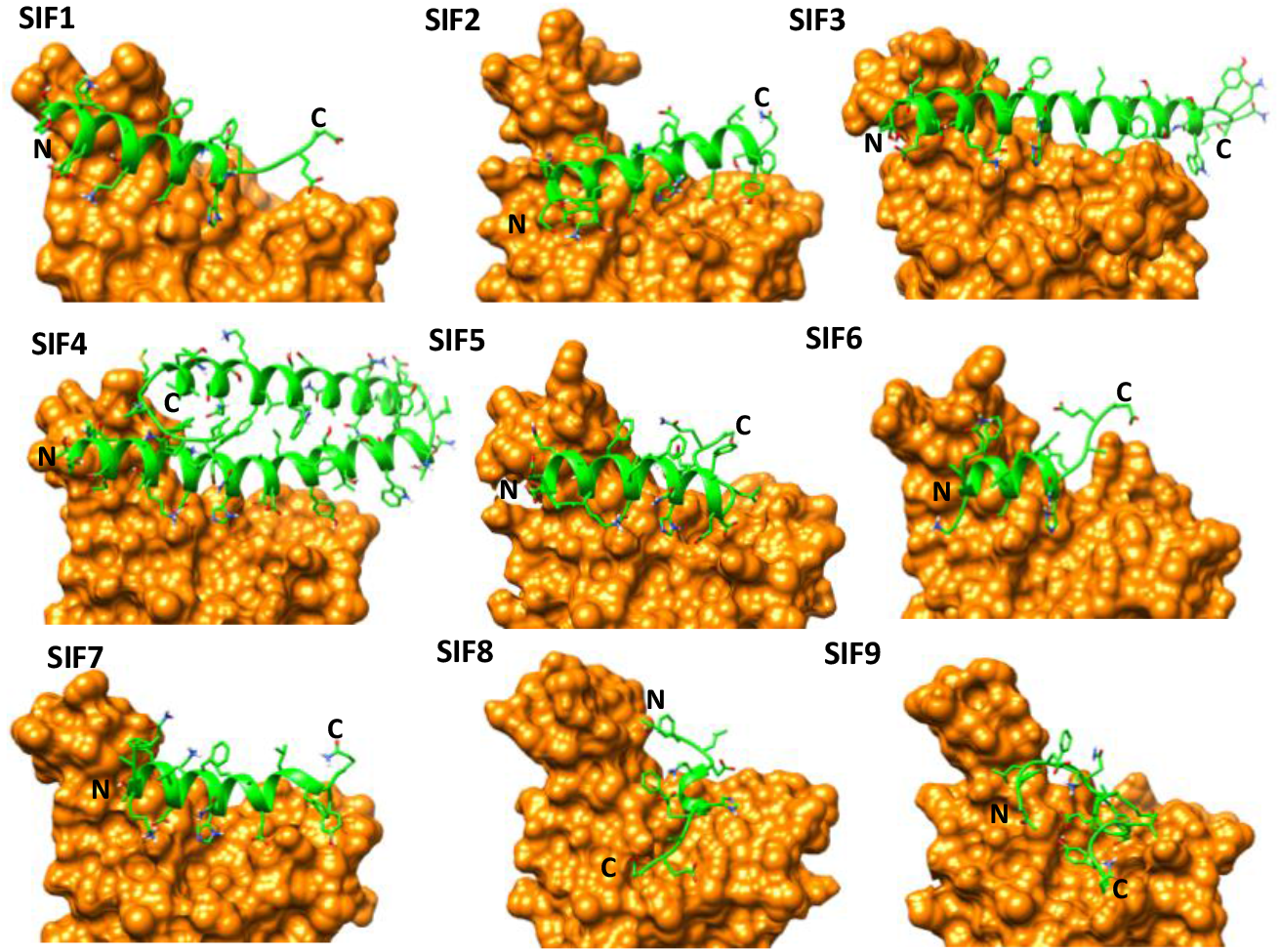
Simulation of the peptides in complex with the RBD of the SARS-CoV-2 spike protein. The structures at the final frame of 300 ns simulations are shown for each complex, with the peptides represented in cartoon in green color and the RBD in surface representation colored in orange. N and C denote N-terminal and C-terminal respectively.

The SIF stability was also measured with their helicity contents in the free and bound forms (**Table 2**). The peptides SIF1 to SIF4 showed high helical contents (~80%) when they are in complex with the spike protein RBD. It is interesting to observe that longer peptides tend to maintain stable helix conformations (**Figure 5**). For example, the two longest peptides, SIF3 and SIF4, are stable in helical conformations in solution as well as bound to RBD. The SIF1 and SIF2 showed enhanced helical conformations when bound to spike RBD, making them potential candidates for peptide drug design.

**Figure 5.**
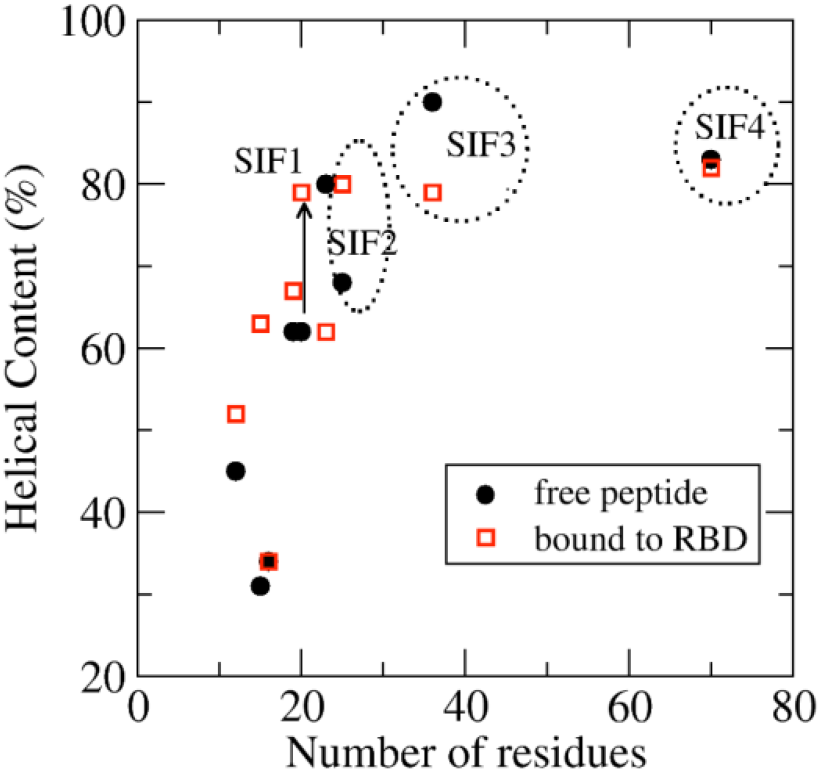
The helical contents of the designed peptides. The SIF3 and SIF4 maintain high helical contents in both free and bound forms. The SIF1 and SIF2 can also be potential candidates for inhibiting peptides.

### Interactions between ACE2 N-terminal fragments and the spike protein RBD open a door for the design of peptidomimetics

MD simulations provided important information about the critical interactions between peptide fragments and RBD. As discussed earlier, residues in RBD interacts with the human ACE2 N-terminal domain mainly via hydrogen bonds and vdW contacts. Compared to the full hACE2, the SIF3 showed stronger binding affinity, reflected in the lower binding energy (**Table 3**). The SIF5 reported in the previous study appeared as the third strongest binder to the RBD, after the SIF4. The free energy calculated from the 300 ns MD simulation data of the SIF5/RBD complex is consistent with the previously reported binding affinity of about 47nM for this peptide^21^. Furthermore, the same study found that the SIF8 did not show binding activities, which can be explained as the unstable helical conformation of the SIF8, which becomes the coiled conformation in solution (**Figure 3**). Even started with the helical conformation in the bound form with the SARS-CoV-2 RBD, the SIF-8 became unfolded and dissociated from the binding pocket of the RBD (**Figure 6**). Combined with experimental data, the simulations not only provide molecular explanations to observations, but also verified the design principles for the hACE2 derived peptides, which are good candidates for binding to spike proteins if the original secondary structures of full hACE2 can be preserved.

**Table 3.** MM-GBSA binding energies of hACE2 and hACE2 derived peptides.

| hACE2 and SIF peptides | MM-GBSA (kcal/mol)# |
| --- | --- |
| hACE2 (Crystal) | -74.65 |
| hACE2 (MD simulation) | -85.88±13.58 |
| SIF1 (19-38) | -49.40±8.07 |
| SIF2 (19-43) | -52.27±10.82 |
| SIF3 (19-54) | -90.43±14.16 |
| SIF4 (19-88) | -67.33±18.08 |
| SIF5 (21-43) | -66.48±9.76 |
| SIF6 (24-38) | -36.52±12.59 |
| SIF7 (24-42) | -54.41±13.9 |
| SIF8 (27-38) | -41.10±9.00 |
| SIF9 (27-42) | -48.23±19.71 |

**Figure 6.**
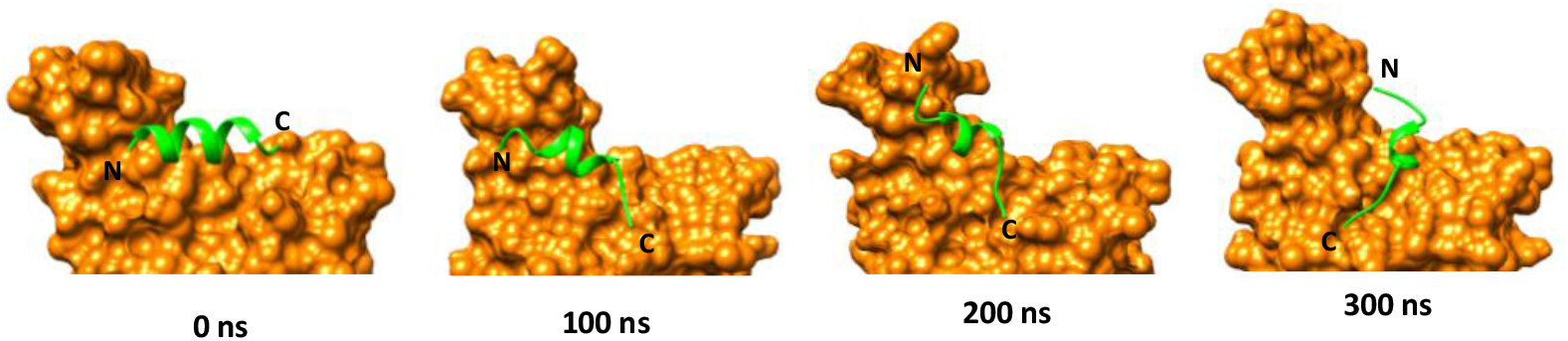
The SIF8 conformational change and the interactions with the Spike protein RBD. The four snapshots from the 300 ns simulation illustrate the conformational changes and the dissociation from the RBD.

## Discussions and Conclusions

It is very encouraging to learn the progress in designing hACE2 derived peptides to inhibit the SARS-CoV-2 spike protein binding to hACE2, such as the hACE2 fragment composed of residues 21-43 (SIF5 in this study) with a disassociation constant (Kd) of 47 nM^21^, and the engineered peptide fragment (based on residues 22-44 and 351-357) that promises higher potency^23^. We have shown that the peptides with the secondary structures as in their hACE2 protein have better chance to be effective in inhibiting the hACE2 binding. **Figure 7** illustrates the binding energy dependency on helical contents, especially the helicity of free peptides (solid black circles in **Figure 7**). Among the strong binders, whose binding energy are lower than −50 kcal/mol, there is a shared sequence segment composed of the residues (24-39). The major difference for SIF’s with stronger binding affinity compared to the common sequence in all SIF’s are the residues of (24-26) and (39-42) (**see Figure 2**), indicating the important roles of these seven residues for the stability and binding affinity for the hACE2 derived peptides. By investigating the properties of free peptides and peptides in complex with the RBD, our approach ensures higher probability of identifying good binders to SARS-CoV-2 RBD. We also demonstrated that the secondary structure preservation and stability in solvent can be probed via MD simulation methods.

**Figure 7.**
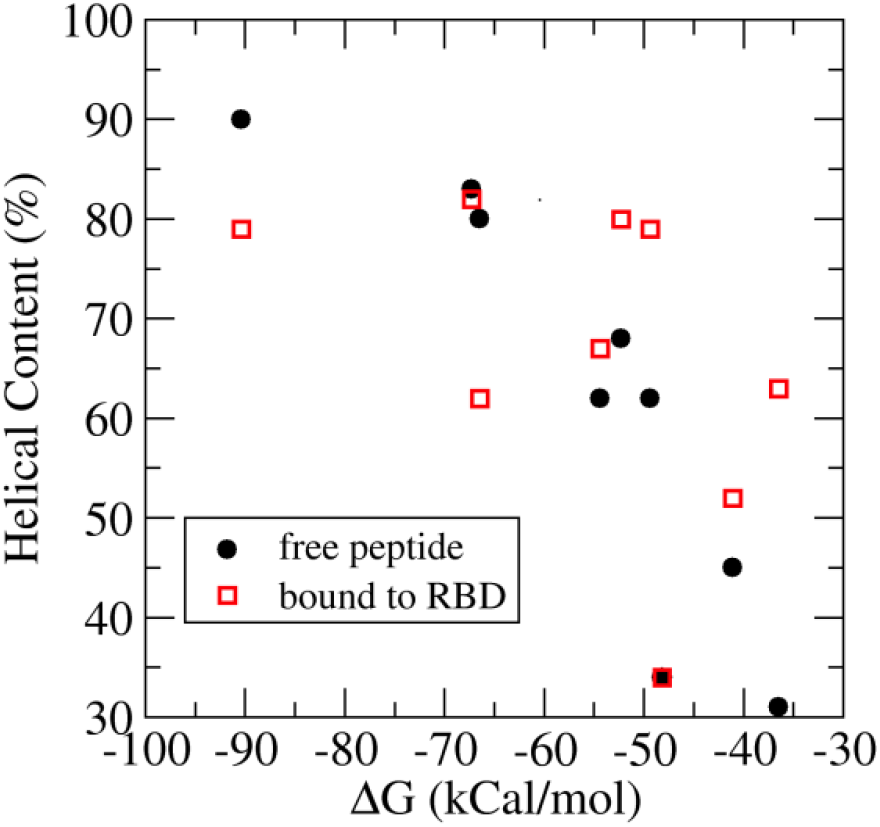
The helical contents and the binding energy are strongly correlated.

SARS-CoV-2 utilizes its spike proteins to gain entry to human cells. The RBD of spike protein is known to interact with the human ACE2 receptor. Hence the disruption of interaction between RBD and ACE2 is an attractive therapeutic option for the treatment of SARS-CoV-2 related disease. In present study, we have identified peptide fragments from the N-terminal region of human ACE2 and investigated their interactions with the spike RBD. MD simulations of peptide fragments with RBD provided insights into important interacting residues at the interface between ACE2 and RBD. We analyzed the conformational stabilities of peptide fragments in water. Three peptide fragments, SIF6, SIF8 and SIF9 appeared to be unstable in water and also showed weak binding with RBD. Among all fragments, the SIF3 showed strongest binding to the RBD. MD simulation data suggest that residues (24-26 and 39-42) of the ACE2 play important roles in the binding of peptides to the RBD. Therefore, these residues should be kept when designing potent peptides. Moreover, binding energies of peptides showed strong correlation with their helical contents in the water. The findings may pave a way for the design of peptidomimetics against SARS-CoV-2.

## Acknowledgement

The work is supported by Beijing Computational Science Research Center (CSRC) via a director discretionary grant. The authors acknowledge the Beijing Super Cloud Computing Center (BSCC) for providing HPC resources that have contributed to the research. The funding from the national natural science foundation (31971136, U1530402) supports the research.

## Competing interests

The authors declare no competing interests.

## Notes

### Competing Interest Statement

The authors have declared no competing interest.

